# An Innovative, Low-Cost Medium for the Bioproduction of Prodigiosin by *Serratia marcescens*

**DOI:** 10.64898/2026.05.07.723488

**Authors:** Lea Massard, Benjamin Toustou, Tiffany Leroy, Alyssia Kassa, Heloise Bauer, Julien Grimaud, Débora Farina Gonçalves

**Affiliations:** Graduate Program in Biotechnology Engineering, SupBiotech, Villejuif, France; Cellule pour la Valorisation de la Recherche Étudiante, SupBiotech, Villejuif, France; Laboratoire CellTechs, SupBiotech, Fontenay-aux-Roses, France

**Keywords:** Prodigiosin, *Serratia marcescens*, solid-state fermentation, cost-effective, HPLC, mass spectrometry

## Abstract

Prodigiosin is a red pigment produced by various bacteria, including *Serratia marcescens*. Despite its wide and promising range of biological activities, the large-scale production of prodigiosin is currently limited by its high cost and low yields. Here we propose and optimize an innovative, low-cost, peanut-based solid culture medium that enhances the yield of prodigiosin produced by *Serratia marcescens*. Colorimetric assays revealed that peanut significantly stimulates prodigiosin synthesis. Further HPLC-MS analysis allowed us to unambiguously identify prodigiosin and shows that our medium specifically improves the yield of prodigiosin. Overall, our innovative culture medium could help lower prodigiosin production costs and, ultimately, open new industrial applications.

## Introduction

Prodigiosin is a tripyrrolic pigment produced by many bacteria, including *Serratia marcescens, Pseudomonas*, and various strains of actinomycetes (Darshan & Manonmani, 2015; Williamson et al., 2006). Since its first identification in 1929 (Wrede & Hettche, 1929), this red pigment has gained attention for its potential use as a natural colorant in food products (Darshan & Manonmani, 2015; Huang et al., 2024). It was not until the 1940s that researchers began to investigate the bioactive properties of prodigiosin (Lichstein & Van De Sand, 1946). Since then, various studies have explored its anticancer potential (Anwar et al., 2022; Darshan & Manonmani, 2015; Ladetto et al., 2025; Manderville, 2001). Other promising bioactive properties have been described, including antimicrobial, antimalarial, antifungal, and immunosuppressive activity (Darshan & Manonmani, 2015; Islan et al., 2022; Ladetto et al., 2025). Overall, prodigiosin has garnered significant interest in the medical sector due to its potential in developing new therapies. Other industrial sectors, including the food and cosmetic industry, have also shown interest in the various bioactive properties of prodigiosin, as this pigment could be used as a natural dye and long-lasting preservative in food and cosmetics (Darshan & Manonmani, 2015; Huang et al., 2024).

In parallel, various studies have focused on the biosynthesis of this pigment (Han et al., 2021; S.-R. Lin et al., 2020; Yip et al., 2019). A member of the Enterobacteriaceae family, *S. marcescens* is a Gram-negative bacterium found in various habitats such as soils, freshwater, wetlands, as well as in human-made environments like bathrooms, sinks, and other damp surfaces (Hejazi & Falkiner, 1997; Ul Huda et al., 2025). In the wild, this bacterium participates in the degradation of organic matter through the production of lipase and chitinase (Hejazi & Falkiner, 1997; Ul Huda et al., 2025). *S. marcescens* is very often harmless to humans but can also be an opportunistic pathogen, responsible for certain nosocomial infections, including respiratory and blood infections (Tavares-Carreon et al., 2023). In biotechnology, *S. marcescens* is prized for its strong synthesis of prodigiosin (Han et al., 2021; S.-R. Lin et al., 2020; Yip et al., 2019).

Despite its numerous bioactive properties and industrial potential, prodigiosin has not yet been exploited commercially. This is due to the current drawbacks of its large-scale production, which presents several major challenges. First, light promotes its rapid degradation and loss of bioactive properties, complicating its storage and applications (Darshan & Manonmani, 2015). Second, despite its relatively high yield compared to other bacteria, the production of prodigiosin by *S. marcescens* remains relatively low, which impairs batch production and increases costs (Han et al., 2021; S.-R. Lin et al., 2020; Yip et al., 2019). Last but not least, current methods of extraction and purification of prodigiosin are complex and expensive, which represents an yet another obstacle to its large-scale use (Paul et al., 2024).

Recent efforts to reduce costs and increase prodigiosin production in *S. marcescens* have mainly focused on optimizing the production of biomass using cost-effective substrates. Researchers have explored the use of agricultural and industrial by-products, such as molasses and glycerol, as alternative, cost-effective carbon sources (Han et al., 2021; Yip et al., 2019). Genetic engineering techniques, notably the overexpression of genes involved in the prodigiosin biosynthesis pathway, have also been tested (Pan et al., 2022; Ul Huda et al., 2025; Williamson et al., 2006). Despite these efforts, large-scale bioproduction of prodigiosin still suffers from the low yields and high costs, which prevents any industrial application. As such, there is a need for more efficient and scalable processes.

One solution lies in the development of innovative, low-cost culture media that would enhance both the growth of *Serratia marcescens* and its prodigiosin synthesis. Peanut is an inexpensive, nutrient-rich compound, which has been proposed as an alternative substrate to optimize prodigiosin production (Bhagwat & Padalia, 2020; Dasgupta Mandal et al., 2021; Giri et al., 2004). This substrate could therefore constitute the basis of a new culture medium.

With the notable exception of one study (Bhagwat & Padalia, 2020), in which the authors screened both liquid and solid media, all previous attempts to grow *S. marcescens* in the presence of peanut products have been conducted in liquid media (Bhagwat & Padalia, 2020; Dasgupta Mandal et al., 2021; Giri et al., 2004; Shekh et al., 2026). While liquid culture offers several advantages, such as scalability, efficient nutrient and oxygen transfer, and compatibility with automation, it also presents notable limitations, including high water and energy demands, increased risk of contamination, complex downstream processing, and potential shear stress on cells (Lizardi-Jiménez & Hernández-Martínez, 2017). Although liquid media remain the dominant choice for industrial-scale bioproduction, solid-state fermentation represents a valuable alternative for certain applications (Lizardi-Jiménez & Hernández-Martínez, 2017). Prodigiosin is hydrophobic (Darshan & Manonmani, 2015; Williamson et al., 2006), a feature that may complicate downstream processing in liquid systems. In addition, solid media are routinely used in research and development (Lizardi-Jiménez & Hernández-Martínez, 2017). This is why we focused our efforts on developing a solid culture medium.

In this study, we propose and optimize an innovative, low-cost, peanut-based solid culture medium that enhances the growth of *Serratia marcescens* and its prodigiosin synthesis. Overall, our culture medium could help lower prodigiosin production costs and, ultimately, allow the adoption of prodigiosin in industrial applications.

## Methods

### Culture Media

The following culture media were prepared:

- Liquid Lysogeny Broth (LB) medium: NaCl (5 g/L), tryptone (10 g/L), and yeast extract (5 g/L) mixed in distilled water.
- Agar LB medium: NaCl (5 g/L), tryptone (10 g/L), yeast extract (5 g/L), agar (15g/L), and sucrose (10g/L) mixed in distilled water.
- Solid peanut culture media, low osmotic stress: liquid LB medium (20g/L), agar (15g/L), sucrose (10g/L), and peanut powder mixed in distilled water. We tested four concentrations of peanut powder: 0g/L (we called this medium M1-NS which stands for “medium 1 – no stress”), 60g/L (M2-NS), 110g/L (M3-NS), and 157g/L (M4-NS). In addition, we prepared one medium with peanut shell (60g/L) instead of peanut powder (M5-NS).
- Solid peanut culture media, high osmotic stress: we added NaCl (15g/L) to media M1-NS, M2-NS, M3-NS, M4-NS, and M5-NS. These new media were called respectively M1-S, M2-S, M3-S, M4-S, and M5-S (the S stands for “stress”).

One may wonder why we did not test peanut powder concentrations above 157 g/L. At this concentration, the medium is thick but still pourable, whereas all attempts to prepare more concentrated media resulted in an almost solid paste that was too viscous to pour into Petri dishes.

All media were autoclaved 20min at 121°C. Solid media were poured into sterile Petri dishes (diameter 90mm, 25mL of medium per dish). All media were then stored at 4°C and used within a week of being prepared.

### Bacterial Culture

Two bacterial strains were used for this study: *S. marcescens* (ATCC 14756) and *Escherichia coli* (DH5α). Frozen stocks of the bacteria were maintained in liquid LB medium supplemented with glycerol (50:50 volume/volume) at −80°C. Precultures were cultivated in liquid LB medium at 28°C with agitation, until they reached an OD_600_ of 1. Solid cultures were prepared by spreading 30μL of preculture onto solid culture plates, in sterile conditions. The plates were then incubated at 28°C for 67h to allow colony development.

### Images Analysis

Two colorimetric quantification methods were used to characterize the color differences observed in the medium. The first method relied on visual assessment by the human eye, while the second employed data analysis using ImageJ software.

For the visual evaluation, a total of 115 participants took part in this assay (age range: 14 to 60 years old; median age: 18; gender: 56 identified as men, 56 as women, 1 as non-binary, and 2 declined to answer). The experiment was conducted over a single day, under constant artificial light, using the same TV screen throughout the test. All participants were seated at approximately the same distance to the screen. This assay was conducted in accordance with all applicable ethical guidelines and regulations for research involving human participants. Informed consent was obtained from all individuals prior to participation. Participants were shown a series of plate pictures, along with a gradient scale (9 rectangles of increasing intensity, going from white to deep red, labeled from 1 to 9) (see Figure 1D). Each participant was asked to rate the perceived intensity of the color on each plate using the gradient scale. The collected responses were aggregated and analyzed as described later in the paper.

**Figure 1.**
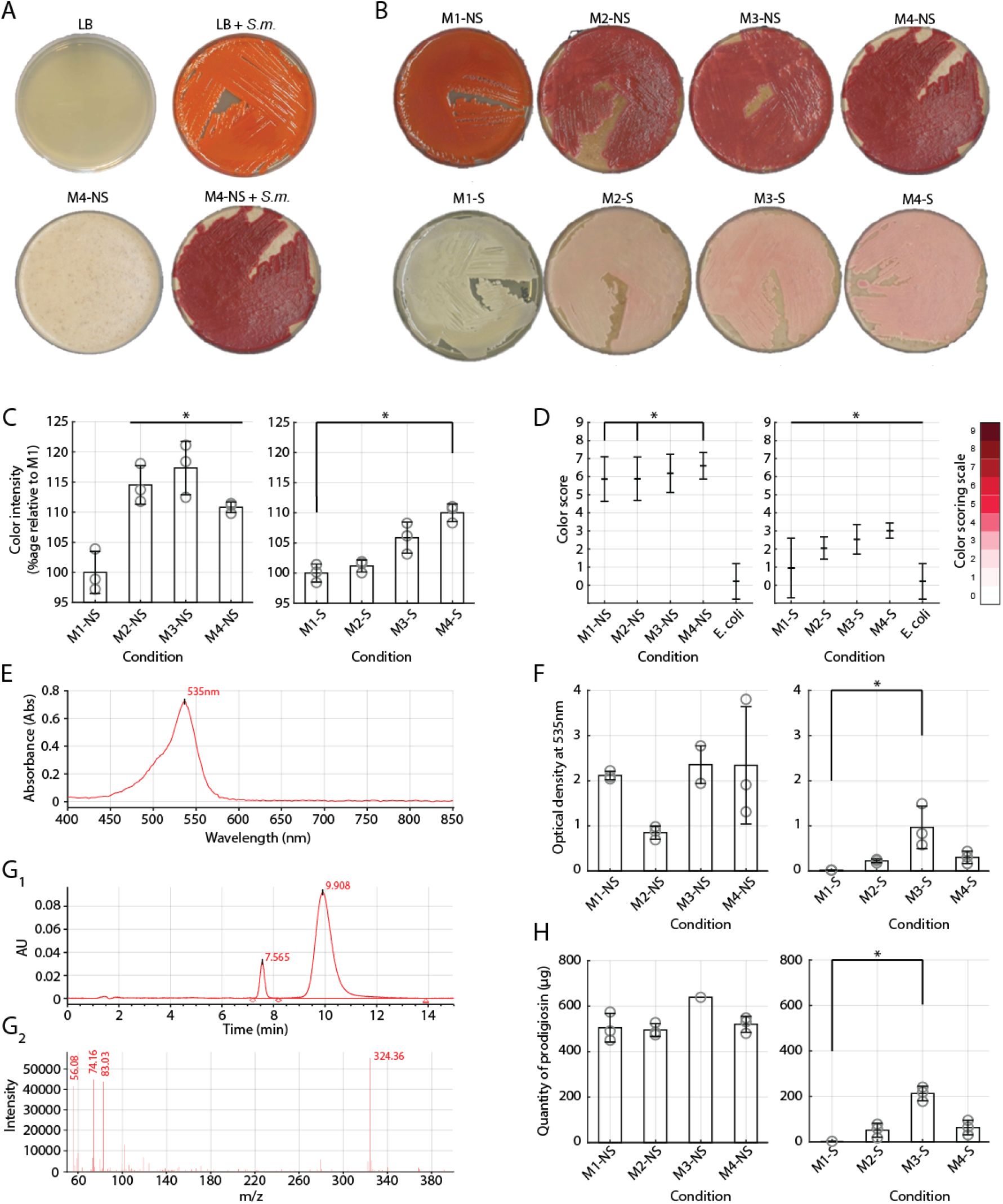
Prodigiosin production by *Serratia marcescens* in different conditions. (A) Upper panel: LB medium alone and LB medium inoculated with *S. marcescens* under non-stress conditions. Lower panel: M4-NS medium (157 g/L peanut powder) alone and M4-NS medium inoculated with *S. marcescens*. (B) Upper panel: Cultures of *S. marcescens* grown in media containing increasing concentrations of peanut powder (M1-NS = 0 g/L, M2-NS = 60 g/L, M3-NS = 110 g/L, M4-NS = 157 g/L) under non-stress conditions. Lower panel: The same media supplemented with NaCl (15 g/L) to induce osmotic stress. (C) Color intensity quantified by ImageJ across the different media. (D) Color intensity evaluated by visual (eye-based) scoring for each medium. (E) Exemplar absorbance spectrum measured after purification. (F) Optical density at 535nm, reflecting relative prodigiosin levels after purification. (G) HPLC-MS chromatographic profiles of prodigiosin. (H) Quantification of prodigiosin concentration obtained from HPLC-MS analysis across the different media. For figures C, F, H quantitative data are presented as mean ± SD. Statistical analysis was performed using a non-parametric one-way ANOVA (Kruskal-Wallis test) followed by Dunnett’s post-test. *p<0,05. Each point represents an individual measurement (n=3 measurements per condition, except panel H condition M3-NS, see Methods). For figure D data are presented as mean ± SD (n=115 responses per condition) and statistical analysis was performed using a non-parametric one-way ANOVA (Kruskal-Wallis test) followed by Tukey’s HSD post-hoc test. *p < 0,05.

For the ImageJ analysis, the mean gray value was used to distinguish differences in color intensity across the selected regions of interest (ROI). A minimum of three independent images were quantified for each condition to ensure reliable measurements.

### Structural Identification of Prodigiosin

#### Prodigiosin Extraction

After incubation, bacteria were collected by thoroughly scraping the surface of the Petri dishes and resuspending in 40mL of ethanol 96%. The ethanol pigment mixture was then frozen at -20°C for 1 hour. After freezing, the tubes were defrosted and centrifuged at 10.000 RPM for 5 minutes at 4°C. The tubes were then frozen again at - 20°C for another hour and subsequently centrifuged under the same conditions (10.000 RPM for 5 minutes at 4°C). The resulting pellet was resuspended in ethanol and vortexed to ensure homogeneity. Sonication was carried out on ice at 35% amplitude for 30 seconds, followed by 30 seconds of rest, repeated for 5 cycles. After sonication, the tubes were centrifuged for 10 minutes at 5.000 RPM at 4°C. The supernatant containing the extracted pigment was carefully collected for further analysis. All supernatants were kept on ice, protected from light by aluminum foil.

#### Purification

After extraction, 1mL of supernatant was filtered on an Amicon 0.2µm membrane after washing with 1 mL of ethanol (EtOH) 96%. The filtrate was completely evaporated at 45°C using a Rotavapor (Darshan & Manonmani, 2015). The resulting dry pellet was resuspended with 100 mL ethyl acetate/water solution (80:20 volume/volume) at room temperature for liquid-liquid extraction. The mixture was transferred into a clean bottle and agitated for 1 hour at room temperature. After mixing, the solution was transferred to a separating funnel and decanted for 1 hour or until complete separation of the two phases (whichever happened first). Two visually distinct phases were obtained: one transparent aqueous phase at the bottom and one red organic phase at the top of the funnel. The aqueous phase was discarded. The organic phase was filtered through a 0.2µm membrane and completely evaporated using a Rotavapor. The final pellet was resuspended in 7 mL of EtOH 96%.

#### Spectrophotometry

Following purification, absorbance was measured with a spectrophotometer (Jenway, model 7315), using standard procedures (de Araújo et al., 2010; Song et al., 2006).

#### High Performance Liquid Chromatography coupled with Mass Spectrometry (HPLC-MS)

After purification, the samples were analysed using HPLC-MS, to identify prodigiosin and evaluate the efficiency of the purification protocol. The analyses were carried out in reverse phase on a Phenomenex Luna C18(2) column (150 mm × 4.6 mm, 5 µm). Chromatographic separation was performed using a Waters Alliance system equipped with a W2690/5 pump, a Waters 2489 UV detector, and an Acquity QDa mass detector (Waters). The mobile phase consisted of a mixture of buffer at pH 4 and acetonitrile (50:50 volume/volume). The flow rate was set at 1mL/min and the column temperature was maintained at 25°C. UV detection was performed at a wavelength of 535nm corresponding to the maximum absorbance of prodigiosin (Williamson et al., 2006). Coupling with a Waters Acquity QDa mass spectrometer was performed in positive ESI mode with a detection range of m/z 50 to 400Da. The ionization parameters included a cone voltage of 15V and a capillary voltage of 0.8kV. These conditions allowed the detection of a signal corresponding to the expected mass of prodigiosin.

#### Statistical Analysis

All visual observations (Figure 1A, 1B) were confirmed on three separate sets of culture media (n=3 observations per condition). Likewise, for Figures 1C, 1F, and 1H, all measurements were performed on three separate sets of culture media (n=3 repeats per condition). Due to technical difficulties during the extraction / purification, we obtained only one exploitable measurement for Figure 1H, condition M3-NS. Finally, for Figure 1D, for each condition, participants were shown pictures from three separate culture media and asked to score based on the three pictures. All 115 participants scored all culture conditions (n=115 responses per condition).

Statistical analyses were performed to compare prodigiosin production across different culture media and experimental conditions. Color intensity quantified by ImageJ, absorbance measurements at 535 nm, and HPLCMS–derived prodigiosin concentrations were analyzed using a nonparametric oneway ANOVA (Kruskal– Wallis test) followed by Dunnett’s posttest for multiple comparisons. Visual (eyebased) color scoring was analyzed using a nonparametric oneway ANOVA (Kruskal–Wallis test) followed by Tukey’s HSD posthoc test. All quantitative data are expressed as mean ± SD, and statistical significance was defined as p < 0,05.

## Results

### Visual Inspection of S. marcescens Growth on Solid Peanut Culture Medium

To document the growth of *S. marcescens* on our new peanut-enriched culture medium, we first conducted a visual inspection before and after inoculation with *S. marcescens*. The formula for our new medium is partly based on classic agar LB medium (see Methods section), which is commonly used for *S. marcescens* culture (Bhagwat & Padalia, 2020; Dasgupta Mandal et al., 2021). We first compared the two media.

Before inoculation, agar LB medium presented a smooth, transparent, uniform, pale golden surface (Figure 1A top-left panel). In contrast, our peanut-enriched medium exhibited an opaque beige texture, slightly firmer to the touch, with small pieces of peanut powder visible throughout the plate (Figure 1A bottom-left panel).

We then compared the appearance of the colonies grown on both media. On the agar LB medium, *S. marcescens* colonies appeared light red, with a smooth, shiny, and slightly creamy texture. Streaking patterns were clearly visible, consistent with localized growth and production of prodigiosin (Figure 1A top-right panel). These observations were consistent with previous reports (Williamson et al., 2006). In comparison, on our peanut-enriched medium, we observed deep red colonies forming a uniform and opaque bacterial mat (Figure 1A bottom-right panel). This mat exhibited slightly domed, dull, and lumpy texture.

Following up on this first set of observations, we asked whether the concentration of peanut powder could impact pigment synthesis. Indeed, *S. marcescens* displayed pigmentation patterns that varied with the concentration of peanut powder (Figure 1B, top plates) As peanut powder concentration increased, enhancement in both pigmentation and colony size were observed. At 157g/L, the highest concentration of peanut powder we tested, the medium was almost entirely covered by a deeply dark red bacterial mat, which shows increased growth and enhanced prodigiosin production, compared to lower concentrations.

We confirmed our visual inspection through two separate colorimetric assays. For the first assay, we quantified red color intensity of bacterial colonies by converting images to grayscale in ImageJ and measuring pixel intensity values within defined colony areas (Figure 1C). For the second assay, we asked 115 participants to grade the intensity of red colonies visible on each plate, using a reference color scoring scale (Figure 1D). Both measurements produced consistent results: intensity scores appeared higher with increasing concentrations of peanut powder, with the deepest red reached for our highest concentration (157g/L) (Figures 1D, left panel). Overall, these results confirm our initial visual inspection (Figure 1B, top plates) and suggest a higher biosynthesis of prodigiosin.

### Analysis of Purified Prodigiosin

We then decided extract and purify the prodigiosin produced on our solid media, with the goal of performing further spectrophotometry and mass spectrometry analyses. In brief, the media were thoroughly scrapped to collect as much biomass as possible. Then prodigiosin was extracted from bacterial cultures using ethanol-based solvent extraction, combined with freeze-thaw cycles, centrifugation, and sonication. The ethanol supernatant containing the pigment was further purified through liquid-liquid extraction with an ethyl acetate/water solution. After phase separation, the red organic phase was isolated, filtered, evaporated, and resuspended in ethanol for subsequent analyses.

Previous studies have reported that the maximum absorbance for prodigiosin was observed around 535 nm (de Araújo et al., 2010; Song et al., 2006). Therefore, we first aimed at using spectrophotometry to confirm our visual inspection. All purified samples showed an absorbance peak at 535nm, consistent with the presence of prodigiosin (Figure 1E). Unfortunately, in our hands, further spectrophotometric quantification showed inconsistencies, including high variability across samples, preventing reliable interpretation (in particular: condition M4-NS) (Figure 1F). This variability may arise from several factors, including slight differences in solvent composition, pH, and unwanted exposure to light, all of which affecting prodigiosin’s spectral properties (Darshan & Manonmani, 2015). Therefore, we did not pursue our initial spectrophotometry strategy.

Instead, to further identify and quantify prodigiosin in the purified sample, we decided to perform an HPLC-MS analysis. HPLC analysis revealed that the primary peak of the purified pigment shared a similar retention time of 10min across all solid media tested, including the standard agar LB medium, suggesting a strong similarity (Figure 1G_1_). Additional peaks with a retention time of 7 to 8min indicated the potential presence of other molecules structurally related to prodigiosin that may have been generated during bioproduction. Furthermore, MS analysis identified a dominant peak corresponding to a mass-to-charge ratio of 324.36 (Figure 1G_2_), which matches that of prodigiosin (Balasubramaniam et al., 2019; Li et al., 2021; Miglani et al., 2023). The MS analysis also revealed the presence of smaller additional peaks (Figure 1G_2_), revealing the presence of prodigiosin analogues, as previously reported (Lee et al., 2011; Li et al., 2021; C. Lin et al., 2019). Nonetheless, based on these combined findings, we identified the red compound observed thus far as prodigiosin.

Further quantification revealed that we purified a similar quantity of pigment (i.e., prodigiosin and the analogues observed in Figure 1G_2_) from all conditions (average ± SD across all samples: 507 ± 40µg per Petri dish) (Figure 1H, left panel). Therefore, it appears that, while the biosynthesis of prodigiosin and analogues is similar in all conditions, our peanut medium specifically enhances the production of red-pigmented prodigiosin (Figure 1B – 1D). In other words, our new medium appears to specifically increase the yield of prodigiosin, with the best results obtained for condition M4-NS.

### Effect of Osmotic Stress on the Synthesis of Prodigiosin

Prodigiosin production by *S. marcescens* can be affected by various environmental stress, in particular osmotic stress. While high concentrations of NaCl (above 40 to 50g/L) appear to impair prodigiosin synthesis (Das et al., 2018; Silverman & Munoz, 1973), concentrations below 40g/L could increase prodigiosin production (Rjazantseva et al., 1994; Tejada et al., 2024). Therefore, we wondered whether prodigiosin production on our peanut-based medium could be improved by a modest addition of NaCl (15g/L).

Upon visual inspection, we found that colonies growing on our peanut-based medium with 15g/L NaCl developed a pink hue, suggesting modest production of prodigiosin (Figure 1B bottom-right panel). These first observations were confirmed through the colorimetric assay described above, as all media containing NaCl were graded as paler than their no-salt counterparts (Figure 1C – 1F). Using the same HPLC-MS approach as above, the retention time and mass-to-charge ratio of the pink pigment confirmed that it was prodigiosin (Figure 1G). Further quantification indicated that the quantity of purified prodigiosin was consistently lower in the media with salt, compared to the media with no salt added (Figure 1H).

Therefore, contrary to what previous reports suggest (Rjazantseva et al., 1994; Tejada et al., 2024), a modest addition of NaCl in our medium did not improve prodigiosin production.

## Discussion

### An Innovative Culture Medium

Here, we developed and optimized a peanut-based solid culture medium for the bioproduction of prodigiosin by *S. marcescens*. Our results demonstrate that this medium supports robust bacterial growth and the yield of active prodigiosin compared to conventional solid media. These findings are consistent with earlier studies that used peanut-derived products as a carbon source in liquid cultures (Bhagwat & Padalia, 2020; Dasgupta Mandal et al., 2021; Giri et al., 2004). Nonetheless, our work is among the first to show that peanut-based solid media can serve as an efficient and cost-effective platform for prodigiosin biosynthesis.

Previous studies have explored other inexpensive substrates for the production of prodigiosin, including glycerol, molasses, and agro-industrial waste products (Kurbanoglu et al., 2015). While these substrates have shown promising results in liquid cultures, our results suggest that peanut-based solid media may provide satisfactory yields without the complexity of liquid culture.

Peanuts are particularly advantageous due to their wide availability, high energy density, and rich nutritional profile. The lipids in peanuts may also serve as important precursors for prodigiosin biosynthesis or may enhance pigment retention in the biomass (Bhagwat & Padalia, 2020; Dasgupta Mandal et al., 2021; Giri et al., 2004). Through which mechanism(s) our enhanced medium leads to an increased yield of active prodigiosin is up for testing.

### A Promising Alternative for Large-Scale Bioproduction

In industry, solid-state fermentation is commonly used to produce enzymes and secondary metabolites, when low-cost production and simpler infrastructure are key (Lizardi-Jiménez & Hernández-Martínez, 2017). Although it is less scalable than liquid fermentation in some contexts, solid-state fermentation can be particularly valuable for high-value, low-volume compounds like prodigiosin. Our peanut-based solid medium could represent a sustainable alternative for small-to medium-scale prodigiosin production, potentially supporting early-stage commercial development.

In addition, our solid medium appears to favor the accumulation of prodigiosin in the biomass, likely due to its hydrophobic properties. This aligns with the known lipophilic nature of prodigiosin, which tends to associate with cell membranes and hydrophobic environments (Williamson et al., 2006). These results suggest that solid-state fermentation may offer advantages for the production and recovery of prodigiosin.

### Towards Low-Cost Bioproduction

One of the major barriers to commercial exploitation of prodigiosin is the high cost of production, particularly due to low yields and expensive downstream processing (de Araújo et al., 2010; C. Lin et al., 2019; Tunca Koyun et al., 2022). Our peanut-based solid medium addresses several of these challenges by providing a nutrient-rich, low-cost substrate that enhances pigment yield and potentially simplifies extraction.

If scaled successfully, this system could lower production costs and facilitate the application of prodigiosin in industries. In biomedical contexts, increased availability of prodigiosin could support its use as a candidate anticancer or antimicrobial agent (Anwar et al., 2022; Arivuselvam et al., 2023; Hejazi & Falkiner, 1997; Islan et al., 2022). In food and cosmetics, its natural pigment properties and bioactivity may offer dual functionality as a dye and preservative (Arivizhivendhan et al., 2018; Darshan & Manonmani, 2015).

### Limitations and Considerations for Scale-Up

Despite these promising findings, our approach bears several limitations. First, the nutritional composition of the medium may vary, depending on the source and processing method of the peanut powder. This variability may affect reproducibility and scale-up. Further testing is necessary to explore this variability and how it may affect yield. Second, while solid-state fermentation offers advantages for small-scale or niche production (Lizardi-Jiménez & Hernández-Martínez, 2017), scaling it up remains technically challenging due to the difficulty of automating solid culture systems. Additionally, we did not address the long-term stability of prodigiosin synthesized on solid media, nor did we explore the efficiency of downstream extraction and purification at larger scales. Further work will be required to evaluate the consistency and robustness of this process under industrially relevant conditions.

Overall, further research should focus on refining and scaling up the peanut-based solid culture system. This includes optimizing the physical parameters of the culture environment, standardizing peanut substrate preparation, and improving downstream processing methods. Techniques such as gradient reverse-phase chromatography, affinity systems, or hydrophobic filters could significantly improve the purity and yield of prodigiosin extracts (de Araújo et al., 2010; Kurbanoglu et al., 2015). The integration of genetic engineering approaches, such as the overexpression of key biosynthetic genes or deletion of metabolic bottlenecks, could further enhance yields when combined with optimized media (Pan et al., 2021, 2022).

## Author Contributions

Project supervision: J.G. and D.F.G. Conceptualization: All authors. Writing - original draft preparation: All authors. Writing - review and editing: All authors. Project administration: J.G.

## Acknowledgements

The authors would like to thank Nedra Liamini for her technical help at various steps of the project, as well as Nassim Arouche for his discussions of the experimental setup and results. In addition, the authors would like to thank Jade Foret, Marina Gayat, and Remy Hubert, who contributed to this project at early stages. The authors would also like to thank Lilian Basset, Kenza Belbaraka, Marianne Chailley, Inès Coquisart, Julien Faure, Jeanne Forest, and Valentine Leroy, for their technical support with regards to the spectrophotometry measurements. Finally, the authors would like to thank Julien Pouard, from ETSL laboratory, for his decisive technical support on the HPLC-MS experiments.

## Funding

This work was supported by SupBiotech’s Research and Teaching Departments.

## Conflict of Interest

The authors declare that the research was conducted in the absence of any commercial or financial relationships that could be construed as a potential conflict of interest.

## Notes

### Competing Interest Statement

The authors have declared no competing interest.

